# Chirp analyzer for estimating amplitude and latency of steady-state auditory envelope following responses

**DOI:** 10.1101/700054

**Authors:** Eduardo Martínez-Montes, Yalina García-Puente, Matías Zañartu, Pavel Prado-Gutiérrez

## Abstract

The envelope following response (EFR) is a scalp-recorded evoked potential elicited by carrier tones or noise, modulated in amplitude with a continuous sweep of modulation frequencies. This non-stationary response reflects the phase-locked neural activity of the auditory pathway to the temporal envelope of sounds and has been commonly assessed by fixed-frequency methods based on the discrete Fourier transform, such as the Fourier Analyzer (FA). In this work, we study the estimation of the EFR with the use of explicit time-frequency methods, which offer more information about the energy distribution of the recorded signal, such as the Short-Term Fourier Transform (STFT) and the Morlet Continuous Wavelet Transform (CWT). We further introduce the Chirp Analyzer (CA), which is similar to FA, but using as basis function the same linear chirp that amplitude-modulates the carrier stimulus. In a direct comparison using controlled simulated responses, the CA showed to be able to estimate the correct EFR amplitudes, without the typical bias offered by the estimation using STFT (equivalent to FA) and more robust to noise than the CWT method, although with higher sensitivity to the presence of a delay in the response with respect to the stimulus. For addressing the latter issue, we also propose here a novel methodology for estimating the apparent latency of the response. This method proved to be reliable when using the STFT and the CA methods, as assessed using simulated responses. The estimation of the EFR amplitude with any of the methods, but especially with CA, should be corrected by using the estimated delay when possible. An illustrative application of these methods to small datasets of a rat and a human newborn, suggested that all time-frequency methods can be used to study the EFR amplitudes in a wide range of modulation frequencies, but they should be interpreted in the light of the limitations shown in the simulation studies.

## I. Introduction

The envelope following response (EFR) is a scalp-recorded evoked potential that reflects the phase-locked neural activity of the auditory pathway to the temporal envelope of sounds (Artieda et al. 2004; Purcell et al. 2004). This auditory response is elicited by non-stationary stimuli, consisting of carrier noise or tones, modulated in amplitude with a continuous sweep of modulation frequencies. Furthermore, the EFR can be evoked by ecologically relevant sounds such as synthetic and natural vowels, reflecting the auditory processing of acoustic features of speech sounds (Aiken and Picton 2006).

The EFR could be clinically valuable for objectively evaluate the outcome of hearing aids, optimizing the fitting process of the hearing devises (Aiken and Picton 2006). Furthermore, the EFR could be potentially used as an objective measurement of temporal auditory processing disorders (Laroche et al. 2013) and learning disabilities. Some of the advantages of evaluation tools based on the acquisition of the EFR are the possibility of eliciting auditory responses using natural stimuli (running speech or stimuli with similar spectral components), the implementation of hearing tests relatively short, and the potential of detecting neural response using statistical tests (Aiken & Picton 2006; Choi et al. 2013). However, a necessary step for the extended clinical application of the EFR is to standardize the procedures for the acquisition and analysis of the response. Studies comparing the accuracy of the methods used to estimate this kind of evoked potential are not available and their limits of applicability need to be established.

Different methodologies have been implemented to analyze the amplitude and phase of the EFR, with the most common method being the Fourier Analyzer (FA) (Purcell et al. 2004; Purcell and John 2010; Prado-Gutiérrez et al. 2012; Choi et al. 2013). Other strategies based on fixed frequency transforms (e.g. the discrete Fourier transform) has also been implemented (Laroche et al. 2013). However, the non-stationary nature of the dynamics of the temporal envelopes of natural sounds, including speech, makes reasonable to use time-frequency methods for characterizing the EFR (Gurtubay et al. 2001; Artieda et al. 2004; Dajani et al. 2005; Perez-Alcazar et al. 2008). Two of the most popular and practical time-frequency methods are the Short Time Fourier Transform (STFT) and the Continuous Wavelet Transform (CWT) using Morlet complex functions (Kiebel et al. 2005). In this context, the Fourier Analyzer (FA) can be seen as a faster variant of the STFT in which the reference sinusoids track the instantaneous frequency of the amplitude envelope of the stimulus instead of estimating the Fourier coefficients for the whole frequency range (Regan 1989).

The variability in the methodology for the analysis of the EFR is not limited to the estimation of the amplitude but extends to the latency of this auditory response, a parameter that can be critical to establishing the neural generators of the evoked potential. Due to the lack of a precise definition of the latency due to its dependence on the method used to estimate it, analyses of oscillatory auditory evoked responses have previously assumed a zero-delay response with respect to stimulus onset (Picton et al. 2002). This might lead, at least, to a miss-estimation of the amplitude of these responses, especially in the case of the EFR. The latency of this response has been calculated as the time shift that maximizes the linear statistical correlation between the envelopes of the stimulus signal and of the electrophysiological response (Laroche et al. 2013). However, in most of the cases, the apparent latency of the EFR is estimated based on the expected linear relationship between the modulation frequency of the stimulus amplitude and the phase difference between the stimulus and the response, as determined by using the Fast Fourier Transform (FFT) and the FA (Kuwada et al. 2002; Purcell et al. 2004; Pauli-Magnus et al. 2007; Prado-Gutierrez et al. 2012). However, these methods are all based on the assumption of stationarity of the electrophysiological signal and to our knowledge, there have not been attempts to estimate the latency for intrinsically non-stationary signals like the EFR.

In this work, we aim to contrast the relative accuracy of time-frequency methods based on the STFT and the CWT, for the analysis of the EFR, stressing their advantages and drawbacks in different scenarios. Furthermore, we introduce the Chirp Analyzer (CA) methodology as a new tool for the reliable estimation of this kind of response. The rationale of the CA is similar to that of the Fourier Analyzer (FA) (Regan 1989; Purcell et al. 2004). The key feature is that CA uses the same non-stationary modulation signals of the stimuli (e.g. chirps) as the reference function, instead of the classical Fourier basis. This follows the hypothesis that the response will match the amplitude envelope of the stimulus and the use of this signal as the reference function will lead to a more precise estimation of the amplitude and phase of the physiological response. We compare the relative robustness of this method with that of the time-frequency methodologies. In this sense, the traditional FA is implicitly included in the comparison, as it can be derived from the STFT. Finally, we evaluate a new methodology to estimate the latency of the EFR, based on measuring the time interval between the onset of every (instantaneous) modulation frequency and the maximum amplitude of the estimated response in the corresponding frequency. For these purposes, we first use simulated data to validate the methods and then illustrate their performance on real auditory responses of human and animal models.

## II. Methods

### A. Theoretical model of the EFR

When the EFR is elicited by acoustic stimuli other than speech-related sounds, the stimulus generally consists in a carrier tone or broadband noise (Fig. 1a) modulated in amplitude by a chirp (Fig. 1b), i.e., a sinusoidal function with a continuous sweep of modulation frequencies, each of which will be called hereinafter instantaneous modulation frequency (IMF). The resulting signal is shown in Fig. 1c. The IMF varies linearly in time, increasing in the first half of the stimulus and decreasing in the second half, with the same rate (Fig. 1d). If the sweep of IMF is slow enough, it can be assumed that the auditory pathway does not respond to the changes in this parameter (Purcell et al. 2004). Therefore, the amplitude of the EFR obtained for a given IMF can be considered as equivalent to the magnitude of the auditory steady-state response elicited by a carrier modulated in amplitude by a sinusoidal signal with that specific IMF. In other words, the amplitude of the EFR can be seen as a measure of the capability of the auditory system to respond to each particular IMF (Purcell et al. 2004). This implies the necessity of developing methods that can estimate the amplitude of the signal for a particular frequency in a particular time segment. In the following section, we will describe the time-frequency methods that will be used in this work to estimate the EFR amplitude and introduce a new method that will be called Chirp Analyzer.

**Fig. 1.**
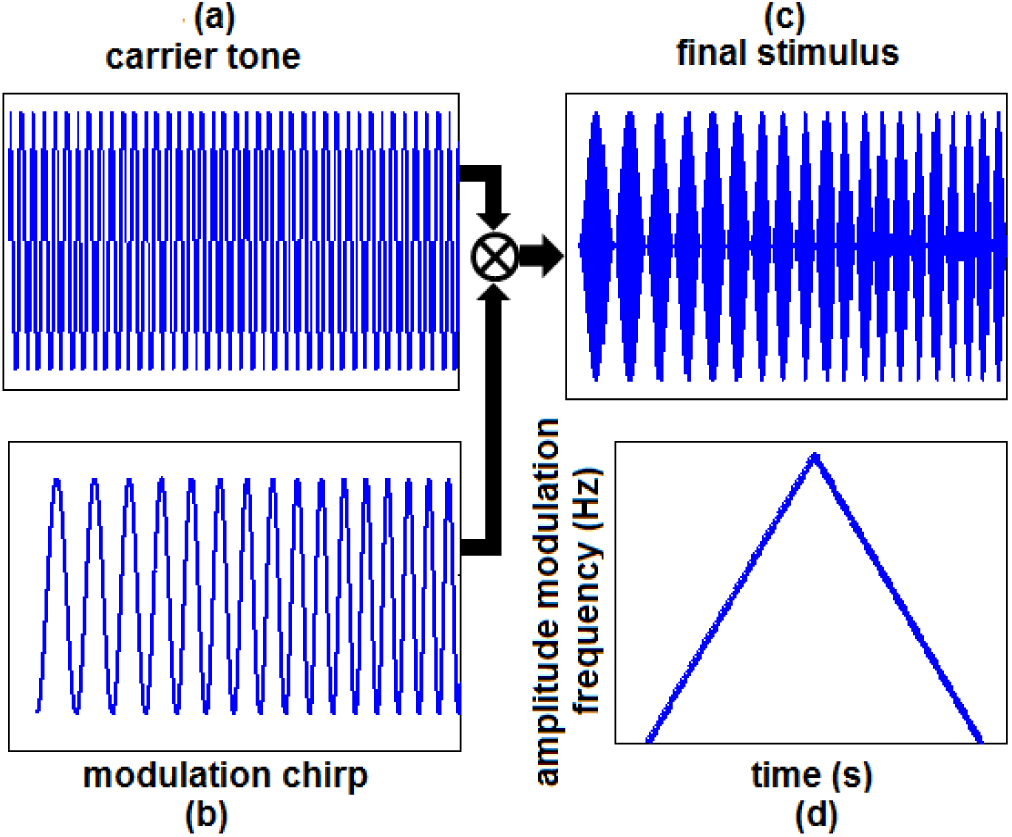
Schematic representation of the first segment of the acoustic stimuli used to evoke the Envelope Following Response (EFR). The amplitude of a carrier stimulus, which can be a pure tone or broadband noise (a) is modulated by a signal with a linearly time-varying instantaneous frequency, known as “linear chirp” (b). A segment of the final stimulus is represented in (c). The total stimulus is composed by a first half where the instantaneous amplitude-modulation frequency increases and a second half where it decreases with the same rate in the same frequency range (d). The latter allows repeating the stimulus continuously without gaps and discontinuities in the instantaneous amplitude-modulation frequency.

### B. Estimating the amplitude of the EFR

#### 1) Short-Time Fourier Transform and Fourier Analyzer

The traditional approach of the Short-Time Fourier Transform (STFT) consists in dividing the non-stationary signal into segments that can be considered stationary. This division can be done by multiplying the signal x(t) by a window function g(t,τ) with a specific temporal width or support (nonzero part of the function) centered at time τ. The STFT is then obtained as the Fourier transform of the product of this window and the signal x(t):

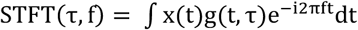

where t and f represent the time points and frequencies, respectively. We used the Goertzel algorithm to estimate the STFT (discrete version) at predetermined frequencies, using a Hamming window of fixed size, which implies constant temporal and spectral resolutions in the whole time-frequency plane (Boashash 2003; Durka 2007). The estimated EFR was extracted as the absolute value of the complex Fourier coefficients in the time-frequency points that correspond to each IMF. By using different window functions, the STFT leads to other methods as particular cases. For example, with the rectangular window function, STFT reduces to the Fourier Analyzer (FA) method described in the literature (Regan 1989; Purcell et al. 2004), which consists in correlating -in time domain-the signal in each temporal window with the sine/cosine reference functions for the IMF corresponding to the center of the window. As an exploratory step, we tested the performance of this method with different window functions and did not find large differences. Nevertheless, the hamming window offered the smallest bias in the estimation of the amplitude. Therefore, in this work, we used the STFT with a Hamming window as a representative of this family of time-frequency methods, including the FA method.

#### 2) Morlet Continuous Wavelet Transform

The continuous wavelet transform (CWT) of signal x(t) is defined by:

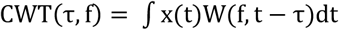

In our case, the function W(f, t) is the complex Morlet “mother wavelet”: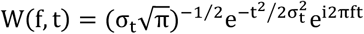, where the temporal support σ_t_ is inversely proportional to the spectral support σ_f_ (Boashash 2003). The magnitude z = f/σ_f_ is kept constant and determines the number of cycles of the wavelet covered in the temporal support (τ ± σ_t_). Therefore, lower spectral resolution and higher temporal resolution are obtained as the IMF (f) increases. That is the reason why this wavelet function is the most popular in EEG analysis (Kiebel et al. 2005; Martínez-Montes et al. 2008). In this study, we used z = 8, which means that the temporal support of every wavelet function will cover 2.5 cycles. With this method, the EFR is then estimated as the absolute values of the wavelet complex coefficients in the time-frequency points corresponding to the instantaneous amplitude-modulation frequencies of the stimulus.

#### 3) Chirp Analyzer

Instead of using a Fourier basis, the Chirp Analyzer (CA) proposed here consists in correlating the signal x(t) with a non-stationary reference function φ(t) that represents the theoretical response, modeled as the analytic (complex-valued) function of the normalized amplitude-modulation chirp used in the stimulus (Fig. 1b):

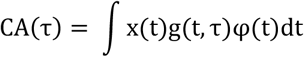

This procedure is carried out in overlapping rectangular windows g(t,τ), for achieving a higher temporal precision with the same spectral resolution. Like in FA, the correlation is performed in the time domain, thus making FA and CA faster than both the STFT and CWT, which need to estimate coefficients for all frequencies at each time point. Correlations with both real and imaginary parts of the reference signal (complex chirps) leads to complex CA coefficients. The EFR is then estimated as the absolute values of the CA complex coefficients in the time points corresponding to each IMF. It is also noticeable that, like in FA, the phase of the complex CA coefficients would reflect the phase of the signal with respect to the reference.

### C. Estimating the latency of the EFR

As mentioned in the Introduction, typical approaches for estimating the latency of oscillatory electrophysiological responses, are based on the estimation of the phase of the signal with the FFT and its variants. As the latency is the delay between two time points, it can be computed by a deterministic calculation or a statistical regression from the known linear relationship between the phase difference (ΔΨ = Ψ_stimulus_ − Ψ_signal recorded_) and the time delay (Δt) in a stationary oscillatory signal with a fixed frequency (f):

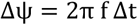

For non-stationary responses such as the EFR, which follows a stimulus-modulating linear chirp with a time-varying instantaneous modulation frequency, IMF, (f(t) = at + b), the phase difference between each time point in the recorded electrophysiological oscillation and the stimulus onset (ΔΨ) will have a quadratic relationship with the time delay (Δt), written in terms of the IMF:

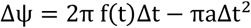

where a is the constant change rate of the IMF. This equation can then be used to estimate the time delay from the roots of the corresponding quadratic polynomial. Alternatively, Eq. 6 can also be seen as a linear relationship between the measured phase differences and the IMFs, allowing the estimation of the time delay from the slope of the corresponding statistical linear regression.

Although the regression procedure for estimating the delay have been used in previous studies based on Fourier analysis (Kuwada et al. 2002; Pauli-Magnus et al. 2007, Prado-Gutierrez et al. 2012), this methodology is highly dependent on the correct measurement of the phases, which has been shown to be unreliable when the level of noise is high, or when the amplitude of the response is small (Sameni & Seraj 2017).

In this work, we propose an alternative approach that takes advantage of the fact that time-frequency methods, such as the STFT and the CWT, give amplitude estimates for all time points in each frequency. This would allow us to look for the maximum amplitude of the response for every specific IMF, in a time interval after the time corresponding to that IMF in the stimulus. We then assume that the time point corresponding to the highest amplitude in that interval will correspond to the latency of the physiological response. An estimate of the neural delay can be obtained as the difference between the time of the IMF in the stimulus and the time of the maximum EFR amplitude for that frequency (see Fig. 2). In the case of the Chirp Analyzer, we compute the response’s amplitude at each IMF by correlating the signal with time-shifted versions of the reference signal (chirp used in the stimulus), in order to find at which time shift we obtain the maximum correlation. This is done only using the real part of the complex chirp, since in this procedure we are not interested in measuring the response’s phase. Similar to the other methods, the delay is then estimated as the difference between the time point of the highest amplitude of the CA coefficients and the time point of the corresponding IMF in the stimulus.

**Fig. 2.**
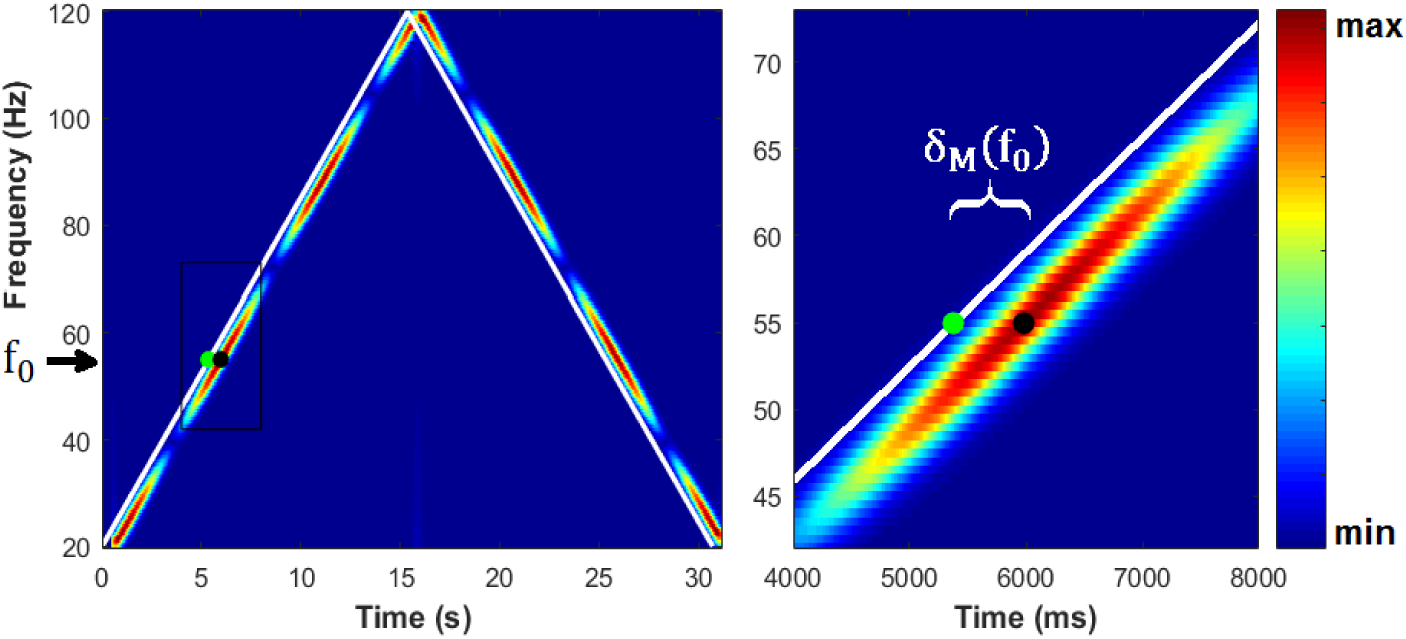
Estimating the delay of the EFR response for each instantaneous modulation frequency f_0_, based on the time-frequency map (left panel). Right panel zooms in the region delimited by a black rectangle in the left panel. White lines represent the (τ_0_, f_0_) points corresponding to every IMF of the stimulus. The neural delay δ_M_(f_0_) for each frequency is estimated by the distance between τ_0_ (green point) and the time at which the highest amplitude of the signal is found (black dot).

Mathematically, for every IMF f_0_, appearing at the time point τ_0_ in the stimulus, the EFR delay is obtained as:

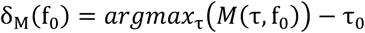

where *M* represents the amplitude of coefficients given by any of the three different methods (STFT, CWT, CA), and the search for the maximum is done for τ ≥ (τ_0_ + 0.2) *s*. If we assume that neural generators of the response to every IMF takes the same time to produce a response, then the EFR delays computed with our method for all IMFs should be the same. Noteworthy, the presence of noise, as well as numerical and model errors of each method, will likely lead to estimate different delays for different IMFs. Therefore, we consider the delays obtained for all different IMFs as random observations of the neural delay. As the statistical properties of these observations are unknown, we evaluate the use of several location statistics such as the mean, the weighted mean (weights proportional to estimated amplitudes), the median, and the mode; as final estimates of the neural delay.

Finally, a very important practical aspect is the definition of the time-frequency precision that will be used for the methods, i.e. the time step and frequency step of results (we avoid using the term resolution, which refers to the capacity of distinguishing different components in the signal due to the basis functions and time windows used in the analysis). For the STFT and CWT, precision is usually linked to the methods but could be controlled by tuning the amount of overlapping of analysis windows. Anyhow, given that the data is usually sampled at a rate much higher than the spectral resolution of these methods, it is convenient to down-sample the resulting coefficients to a smaller amount of (τ, f) pairs. Likewise, the CA can in principle be computed for every time sample (windows overlap in all points but one) but it is easy to change the amount of overlapping to obtain a longer time step (smaller precision). Higher temporal precision would provide a better scenario for estimating latencies but will lead to a higher computational burden. Moreover, higher precision will not lead to better estimates of the EFR, as long as the temporal resolution of the method remains the same. A practical criterion on the more convenient time precision should be followed as a tradeoff between obtaining a reliable delay estimation and a reasonable computational cost. For instance, we here used the highest time precision for estimating delays, but when the delay was known or was not of interest, we used the time step corresponding to the frequency precision imposed for all methods (0.5 Hz), since the estimation of the response in intermediate frequency values will not give us better spectral resolution and the estimated EFR will be practically the same.

### D. Simulated and real datasets

Fig. 3 shows a schematic representation of the model followed to simulate EFR responses, which will allow to evaluate the performance of the methods studied here to estimate these responses. The key point is that we model the electrophysiological response as being the same as the modulation signal of the stimulus (the chirp) but with different instantaneous amplitudes (envelope). Therefore, we can simulate any desired EFR response and use it to modulate the stimulus chirp to create the ideal electrophysiological response (Fig. 3, bottom row), which is then perturbed with noise to emulate typical noisy measurements (Fig. 3, top row).

In order to generate the simulated data, we performed a virtual experiment with a stimulus consisting in a carrier tone (a fixed-frequency sine function), modulated by a chirp signal with an IMF linearly increasing from 20 to 120 Hz in the first half (15.36 s), and decreasing with the same rate in the second half (see scheme in Fig. 1d), for a total duration of 30.72 s. To obtain the corresponding virtual recordings, we simulate a “true” EFR as the instantaneous amplitude for each IMF, which is then multiplied by the modulation chirp. Finally, the resulting signal is added to white Gaussian noise, with zero mean and a variable variance that allows for controlling the SNR of the synthetic “recorded data”.

The robustness of the methods for estimating the EFR was tested by simulating electrophysiological responses with three different shapes with respect to the instantaneous modulation frequencies: 1) *Sine-deep*: the true EFR is simulated as the absolute value of a sinusoid function with 100% of modulation depth, which means that the system does no respond to the stimulus in particular IMFs (where true EFR is zero); 2) *Sine-low*: the true EFR follows a sinusoid function with 50% of modulation depth, leading to nonzero response to every IMF although with smoothly varying magnitudes and 3) *Rect-deep*: the true EFR is simulated as a rectangular function, which models sharp changes in the magnitude of the response and only nonzero responses in particular frequency bands. Fig. 4 shows these simulated “true” EFR in red traces. In all cases, white noise was added with different peak signal-to-noise ratios (pSNR = 2, 1 and 0.1), defined as the ratio between the squared maximum of the signal (fixed to 1 in all simulations) and the variance of the noise. To explore the influence of the latency of the response in the estimated EFR and evaluate the performance of the different strategies for estimating it, we simulated responses with latencies of 10, 50, and 100 ms, to cover a wide range of delays.

**Fig. 4.**
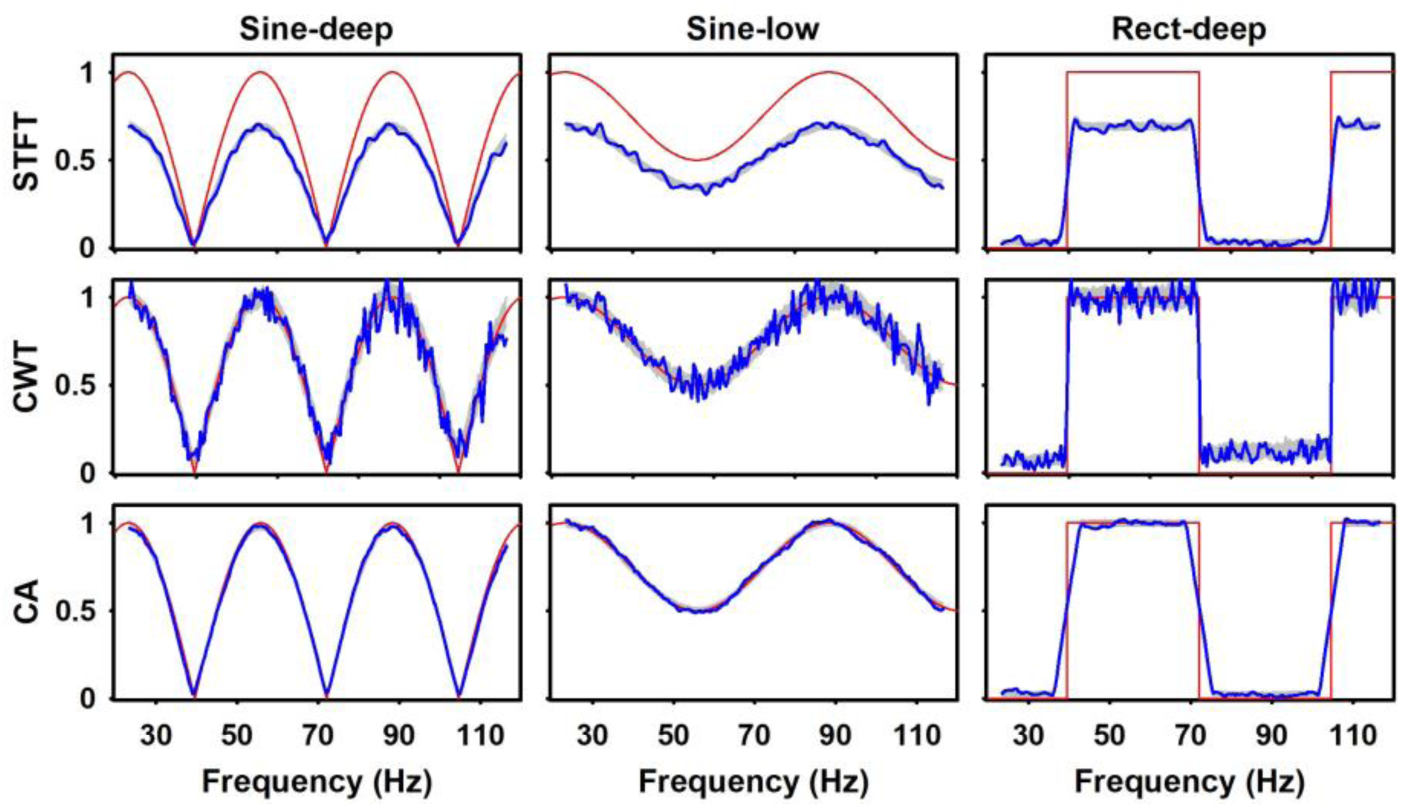
EFR estimations by the three methods (top: STFT; middle: CWT; bottom: CA) from data created with simulated EFR (red lines) of different shapes (left: Sine-deep; central: Sine-low; right: Rect-deep). Blue lines represent the estimated EFRs for a single run, while the grey shadows cover one standard deviation around the mean EFR across the 50 repetitions of each simulation. All the signals were simulated with pSNR=2. The amplitudes are reported in arbitrary units. Note that estimated EFRs do not have values close to the edges, as they are obtained in time windows, such that the first and last values correspond to the time of the center of the first and last complete windows.

In every simulated scenario, we created 50 realizations with different random noise and computed the mean and standard deviation of the results (estimated EFR and/or delays). However, the estimated EFRs are shown for a typical run, together with a shadow covering one standard deviation around the mean across all repetitions. We think this is the correct way to evaluate the performance of the methods since in practice, we can only estimate the EFR from a single signal and not as the mean across many estimations, which usually looks smoother and more robust.

To test the methods in the analysis of real data, we used electrophysiological recordings obtained from adult rats (age: 70 postnatal days) and humans newborns (age range 5-18 days). Devices and methods for data acquisition in each population are detailed in Prado-Gutierrez et al. 2012 and Mijares-Nodarse et al. 2012, respectively. Briefly, the stimulus consisted in a broadband noise as the carrier, modulated in amplitude by a chirp with a continuous sweep of amplitude-modulation frequencies. The IMFs increased linearly from 20 Hz to 200 Hz in the first half and decreased symmetrically in the second, as illustrated in Fig. 1d. Acoustic stimulation was delivered at 50- and 70-dB SPL in the case of rats and only at 50 dB HL for human newborns. In both cases, the stimuli were presented monaurally with ER 3A Etymotic Research insertion earphones. Thirty 30.72-s sweeps of the whole stimulus segment were presented continuously. The 30 segments were synchronized to the stimulation and then averaged in the time domain to increase the SNR. As mentioned in Prado-Gutierrez et al. (2012), the recordings in rats were acquired in anesthetized animals, using a combination of ketamine and diazepam. Since ketamine increases the amplitude of the ASSR to low IMFs in rodents (Kuwada et al, 2002), the EFR amplitude of the response to these IMF (mainly below 70 Hz) was highly variable and not reproducible. In human newborns, the ASSR is most easily and consistently recorded when the amplitude of the acoustic stimulus is modulated at frequencies higher than 60 Hz and 100 Hz (Rickards et al. 1994). Therefore, we restricted the analyses of the EFR to the 90-to 190-Hz IMF range, to be able to compare the results obtained in rats and human newborns.

## III. Results

### A. Analysis of simulated data

#### 1) Estimation of EFR amplitudes

All the methods were able to reproduce the forms of the simulated EFR, although the accuracy of the estimations strongly depended on the methods of analysis (Fig. 4). For instance, the estimated EFRs with both the CWT and the CA were very similar in amplitude to the simulated response, while the EFR estimated with the STFT showed a consistent bias toward lower amplitudes (of about 25% lower than the simulated amplitude), independently of the shape of the simulated signal. Quantitatively, correlations were above 0.90 for all methods in all scenarios, although it was higher for STFT and CA in the sinusoidal responses while higher for the CWT in the rectangular simulation. The latter could be explained by the lower temporal resolution of STFT and CA, which cannot follow the abrupt changes in the rectangular simulation. The relative error (relative Euclidean distance between the estimated and simulated EFRs) were always higher for STFT (above 30%), while again CA was the lowest (below 5%) for the sinusoidal simulations and CWT the lowest (15%) for the rectangular one. Regarding computational time, STFT and CWT are slower (as they need to compute the whole time-frequency map) and strongly depend on the number of frequencies to analyze and the length of the signal. In an Intel Quad-Core i7 CPU, with 16 GB of RAM, they took about 100 s for each run, while the CA took about 20 s in average for each single run.

#### 2) Noise robustness of the estimated EFR

The influence of the level of noise on the estimation of the EFR was analyzed in the case of the Sine-deep simulation (Fig. 5). As expected, the reliability of the EFR estimation was affected when the pSNR decreased, for all methods. However, when the output of the three methods were contrasted in signals with equal pSNR, the CWT was the most sensitive to the presence of noise in the recordings (Fig. 5, middle row). The noise impaired the estimation of the EFR performed by using the STFT in a lesser extent, while the CA was the most robust method to increases in the noise level of the simulations (Fig. 5, top and bottom rows, respectively). The correlation between the estimated EFRs and the simulated ones in this case, were higher than 0.95 for all methods except in the case of very noisy data (pSNR of 0.1), where CA reached 0.97, STFT 0.92 and CWT 0.63, in average across repetitions. The relative error was not affected by the increased of noise level in the case of STFT (30% to 31%, for pSNR=2 to pSNR=0.1, respectively), while increased for CA from 3% to 10% and for CWT from 9% to 46%.

**Fig. 5.**
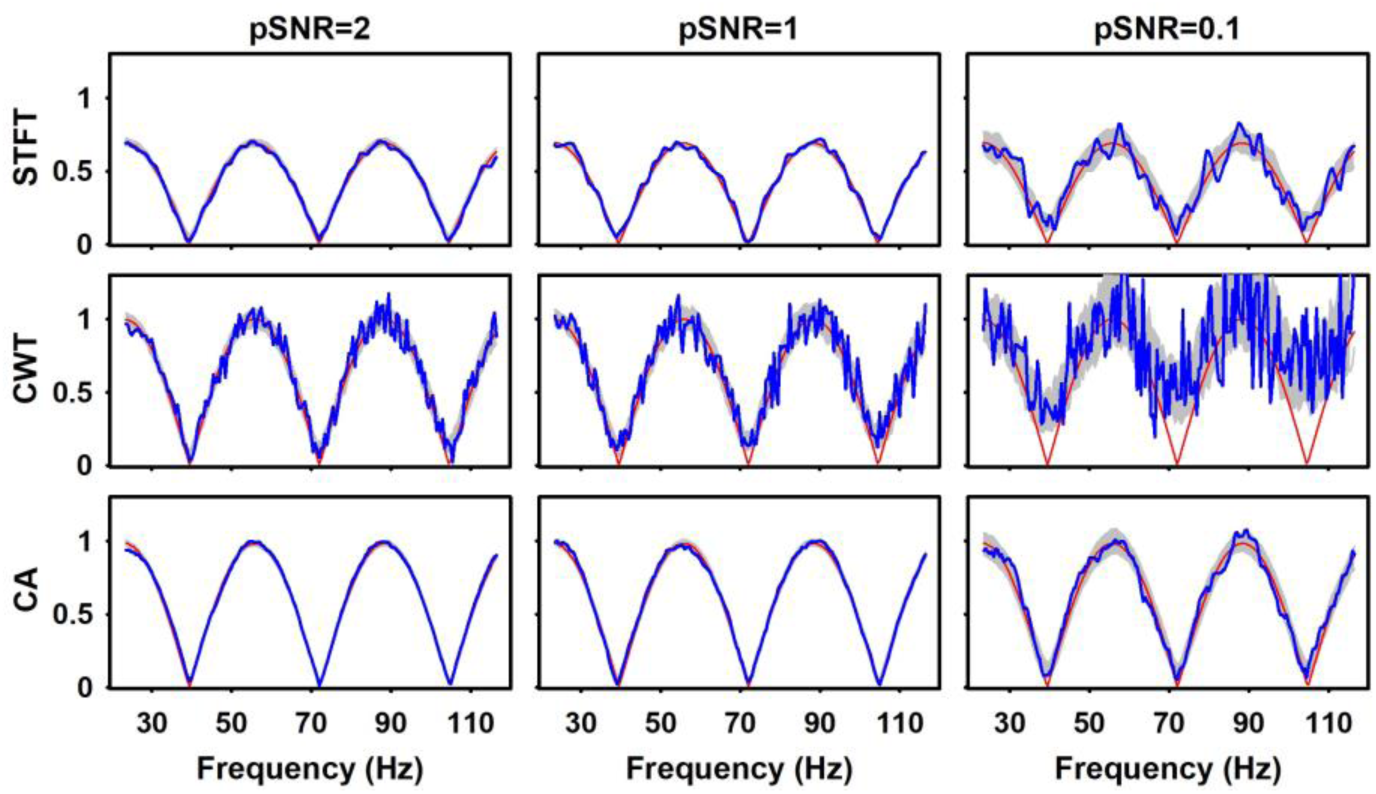
EFRs estimated from data created with simulated Sine-deep EFR, using different levels of noise (pSNR of 2, 1 and 0.1). Blue lines represent the estimated EFRs for a single run, while the grey shadows cover one standard deviation around the mean EFR across the 50 repetitions of each simulation. The red lines represent the EFR estimated when there is no noise affecting the simulated data. The amplitudes are reported in arbitrary units.

#### 3) Influence of the response latency in the estimated EFR

Computation of the EFR amplitude assumes physiological responses with zero delay. Since the latency of neurons in the ascending auditory pathway can be up to dozens of milliseconds, the EFR amplitude might be biased when the latency is not considered in the estimation of the response amplitude. To test this hypothesis, we simulated signals using three different delays (10, 50 and 100 ms) and compared the EFR estimated assuming that there is no delay (i.e. the typical procedure) with the EFR obtained when the mismatch introduced by the delay were corrected -hereinafter called delay-corrected response (Fig. 6).

**Fig. 6.**
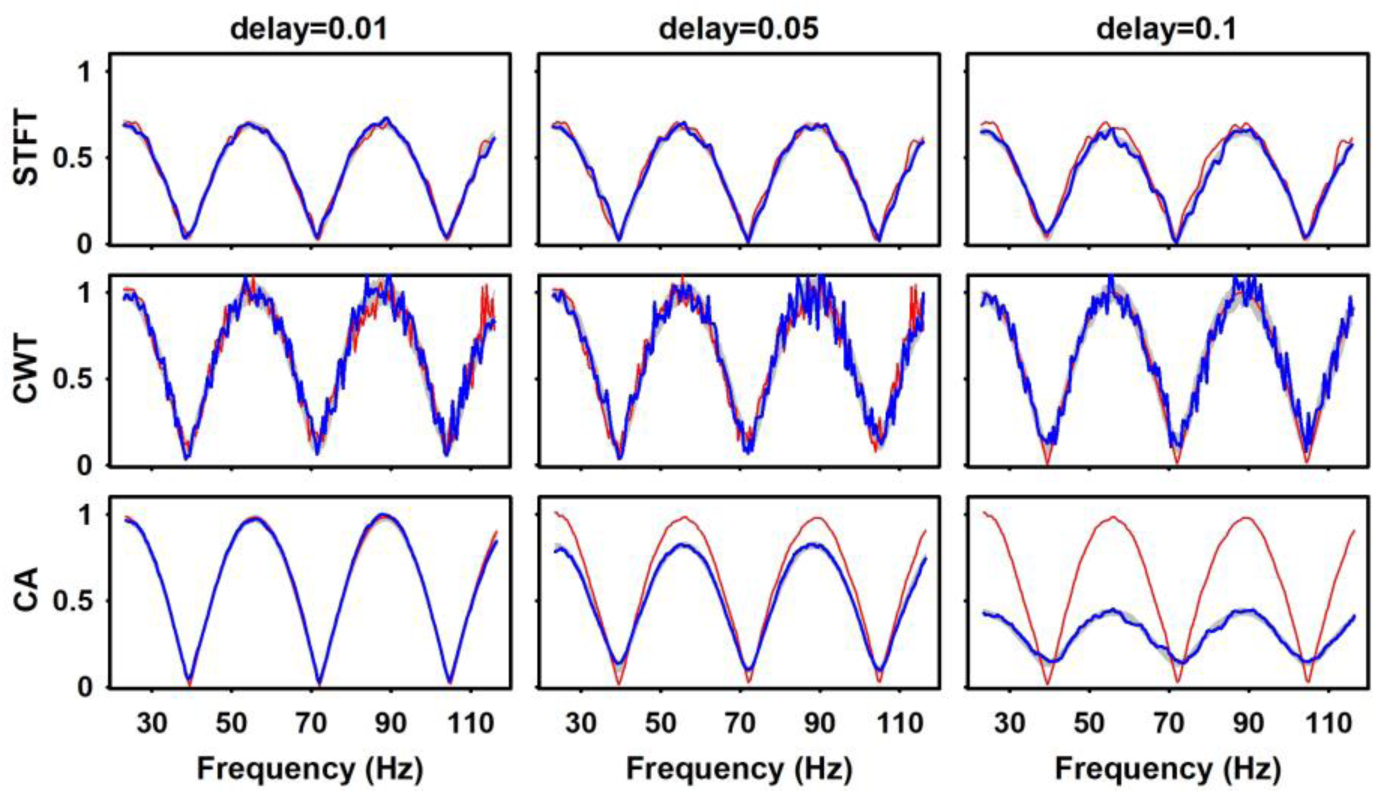
EFRs estimated from data created with simulated Sine-deep EFR, using different delays with respect to the stimulus (10, 50 and 100 ms). Blue lines represent the estimated EFRs for a single run, while the grey shadows cover one standard deviation around the mean EFR across the 50 repetitions of each simulation. The red lines represent the EFR estimated considering the correct delay in the response, i.e. delay-corrected responses. The amplitudes are reported in arbitrary units.

Overall, the delay-corrected EFRs computed with the STFT and CWT matched the corresponding ground truth signal (Fig. 6 upper and middle panels). On the contrary, the amplitude of the EFR computed with CA was systematically underestimated as responses had longer delays, which implies that this method is more sensitive to the presence of a delay between the stimulus and the physiological response. Considering that the stimulus-response mismatch introduced by the acquisition system is usually negligible, these results suggest that analyzing the estimated EFR amplitude for each frequency in a time window after the presentation of the corresponding IMF in the stimulus, might be helpful to estimate the latency of the electrophysiological response, which is the topic addressed in the next section.

#### 4) Estimation of the latency

We calculated the instantaneous delays δ_M_(f_0_) (for the different methods M = {|STFT(τ, f)|; |CWT(τ, f)|; |CA(τ)|}) and computed the following measures of central tendency: weighted mean, mode and median. We also estimate the delay as the slope of a linear regression between estimated phases and instantaneous frequencies. Table I shows, for each of the measures, the mean and standard deviation of absolute errors in ms (absolute value of the difference between the simulated and estimated delays) across the 50 repetitions. This was performed for the Sine-deep simulation scenario, with a true delay of 50 ms and using different levels of background noise.

**TABLE I.**
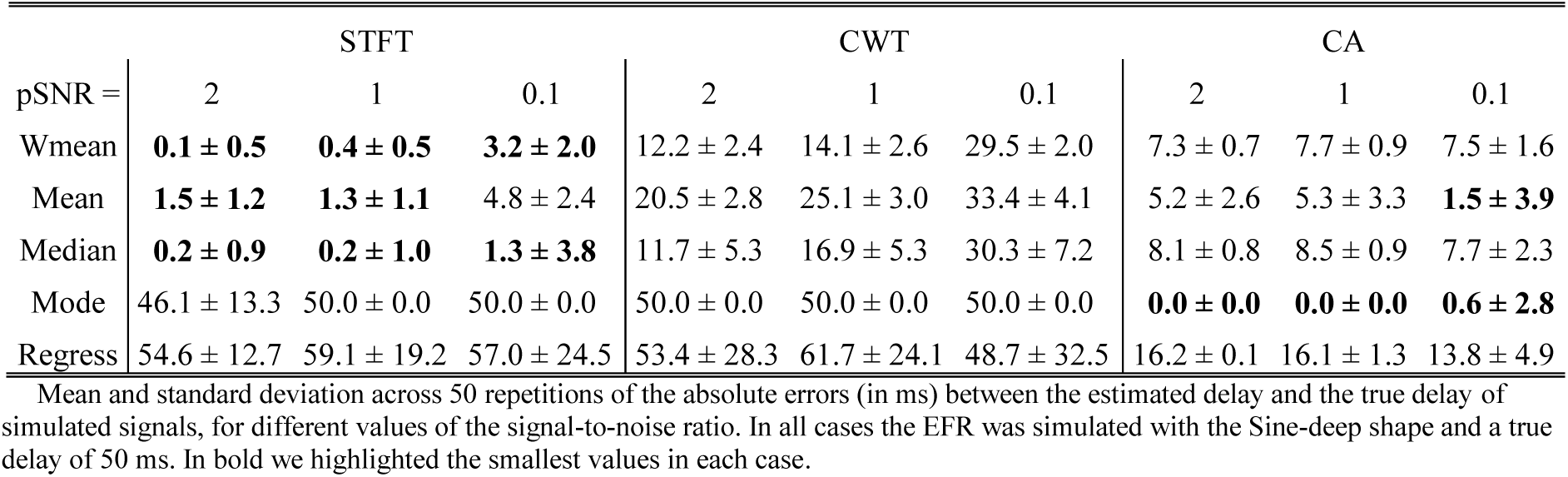
Absolute errors of estimated delays for different noise levels

As expected, the errors typically increased as the SNR of the recording decreased (Table I). Furthermore, the estimation of latencies computed with each method using the regression approach consistently showed larger errors. The EFR latencies estimated using CWT consistently showed higher absolute errors as compared with the other methods. Therefore, the CWT was not considered for further analysis of latency or delay estimation. Using the STFT, the median, weighted mean and mean of all delays estimated for each IMF, offered very good estimates of the true delay, with errors below 2 ms. However, the best results were obtained using the mode of all delays estimated with the CA.

To provide more elements about the statistical measure most suitable for the estimation of the EFR latency, we analyzed simulations with different true delays (Table II). Again, the STFT estimates by using the median, weighted mean, and mean were very accurate in the case or large delays (50 and 100 ms), while better estimates were obtained with CA for smaller delays. However, with this good signal-to-noise ratio (pSNR=2), the mode of the values estimated by the CA was more exact in all cases. The regression approach offered the highest errors for both methods, especially in large true delays.

**TABLE II.** Absolute errors of estimated delays for different true delays

| True delay = | STFT |  |  |  | CA |  |  |  |
| --- | --- | --- | --- | --- | --- | --- | --- | --- |
|  | 0 | 10 | 50 | 100 | 0 | 10 | 50 | 100 |
| Wmean | 8.2 ± 0.4 | 4.2 ± 0.4 | **0.1 ± 0.4** | **2.6 ± 0.5** | **1.3 ± 0.2** | **2.2 ± 0.3** | 7.4 ± 0.8 | 13.4 ± 1.1 |
| Mean | 11.8 ± 1.0 | 7.7 ± 1.3 | **1.2 ± 1.1** | **3.0 ± 1.3** | 9.1 ± 0.8 | 5.6 ± 0.8 | 5.6 ± 2.7 | 17.2 ± 2.1 |
| Median | 3.4 ± 0.5 | **0.5 ± 0.8** | **0.1 ± 0.7** | **0.4 ± 0.8** | **0.0 ± 0.0** | **0.1 ± 0.2** | 8.3 ± 0.8 | 10.2 ± 0.5 |
| Mode | **0.0 ± 0.0** | 10.0 ± 0.0 | 46.1 ± 13.3 | 45.8 ± 50.8 | **0.0 ± 0.0** | **0.0 ± 0.0** | **0.0 ± 0.0** | **0.2 ± 1.2** |
| Regress | 5.7 ± 15.7 | 10.3 ± 16.6 | 52.7 ± 12.2 | 113.2 ± 17.8 | 0.9 ± 9.5 | 10.9 ± 7.5 | 16.2 ± 0.1 | 16.1 ± 0.1 |

Mean and standard deviation across 50 repetitions of the absolute errors (in ms) between the estimated delay and the true delay of simulated signals, for different values of the true delay. In all cases the EFR was simulated with the Sine-deep shape and pSNR=2. In bold we highlighted those values with both mean and std (of absolute errors) not higher than 3 ms.

The reliability of the measures to estimate the latency of the EFR was further tested in simulated recordings with different EFR shapes. Fig. 7 shows the instantaneous delays estimated for each frequency with the STFT and the CA, for the three simulated shapes, a true delay of 50 ms and pSNR = 2. The high standard deviation suggest that estimations of the instantaneous delays were not reliable at those frequencies for which the amplitude of the response was very small. This was mainly evident for the STFT when the amplitude of the EFR was close to the noise floor (std larger than 200 ms), which occurred in the case of Sine-deep and Rect-deep simulations. However, the mean values of all instantaneous delays estimated with the STFT are more accurate than those estimated with the CA, which seems to slightly underestimate the simulated values in all scenarios (black horizontal line). This explains why the STFT offers better estimates using the mean and wmean (see Table I and II, and values above each panel in Fig. 7), while the CA is better using the mode, since the instantaneous delays are randomly biased but the correct value is the most repeated one, although only in a few frequencies.

**Fig. 7.**
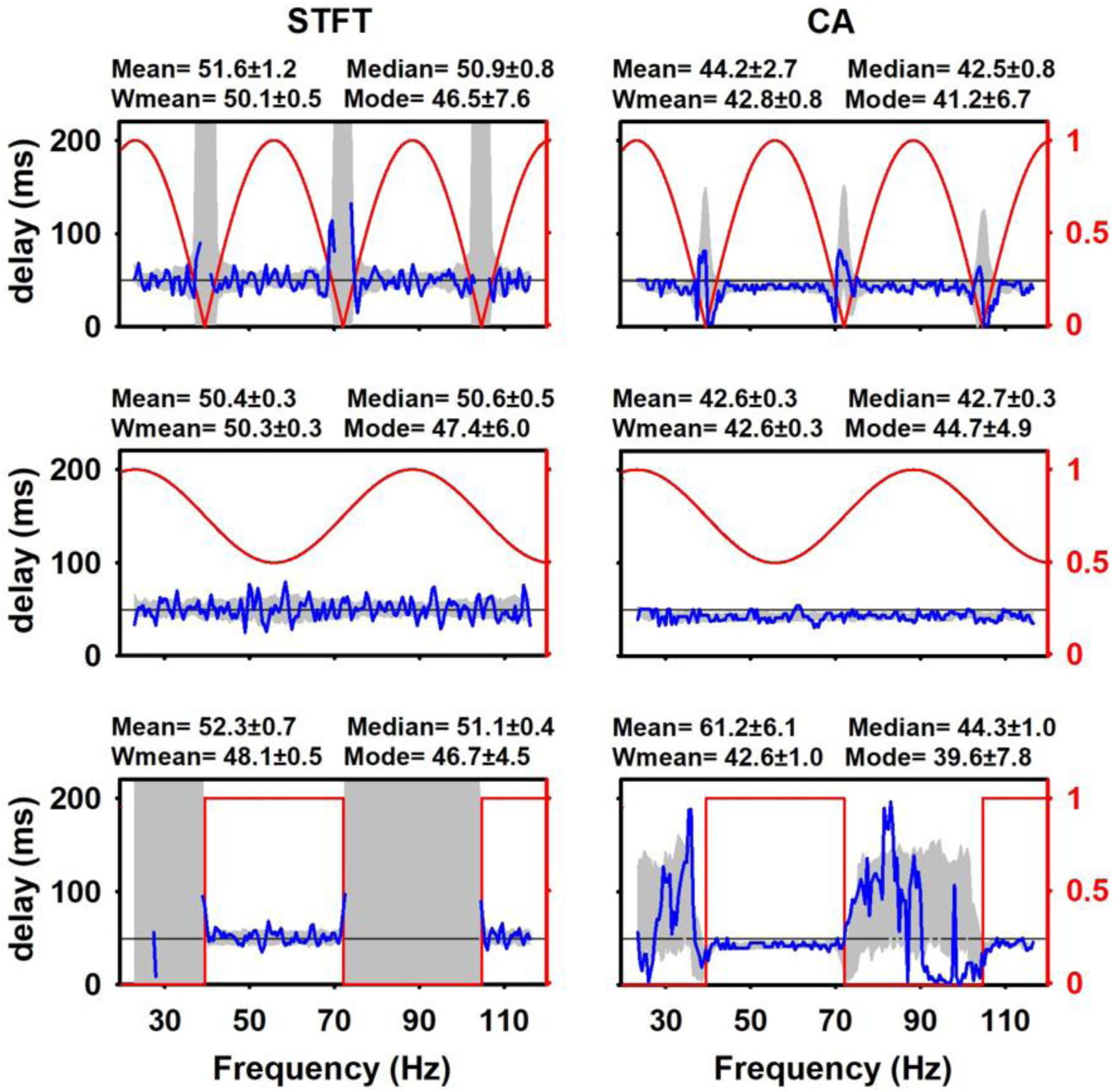
Estimation of instantaneous delays for a single repetition (blue lines) and one standard deviation around the mean across all 50 repetitions (shadow), as estimated by the STFT and the CA methods, from simulations using the different shapes of the true EFR (top row: Sine-deep; middle row: Sine-low; bottom row: Rect-deep). The true EFR in arbitrary units is underlaid with red lines just for illustrative purposes. The signals were simulated with pSNR = 2 and a true latency of 50 ms. In each panel, four values of the estimated latency are given as the average across the 50 repetitions of the weighted mean (Wmean), mean, median and mode of all instantaneous estimated delays.

Given that the CA method is the most sensitive to the presence of a nonzero delay, we explored the practical usefulness of estimating the delay with the methods proposed here. We compared the EFR estimated with the CA without correcting for any delay (i.e. assuming zero delay) and the EFR computed with a correction using the estimated delay. This was studied using three different delays (10 ms, 50 ms, 100 ms) for simulating the response, and for the three shapes in order to evaluate the influence of the EFR amplitudes. The correction was done using the delay estimated as the mode of the instantaneous delays obtained with the CA method (Fig. 8). We found that the estimation of the EFR amplitude was considerably improved in the whole frequency range, even for those frequencies where the actual response’s amplitude was very small or zero. Reconstruction of the EFR was very good in all cases (relative error lower than 0.5% for small delay and less than 2% for delays of 50 and 100 ms). This result was consistent when using the estimated delays with the rest of the population measures (wmean, mean and median), although the estimated and the true delays differed in a range of 5 to 10 ms.

**Fig. 8.**
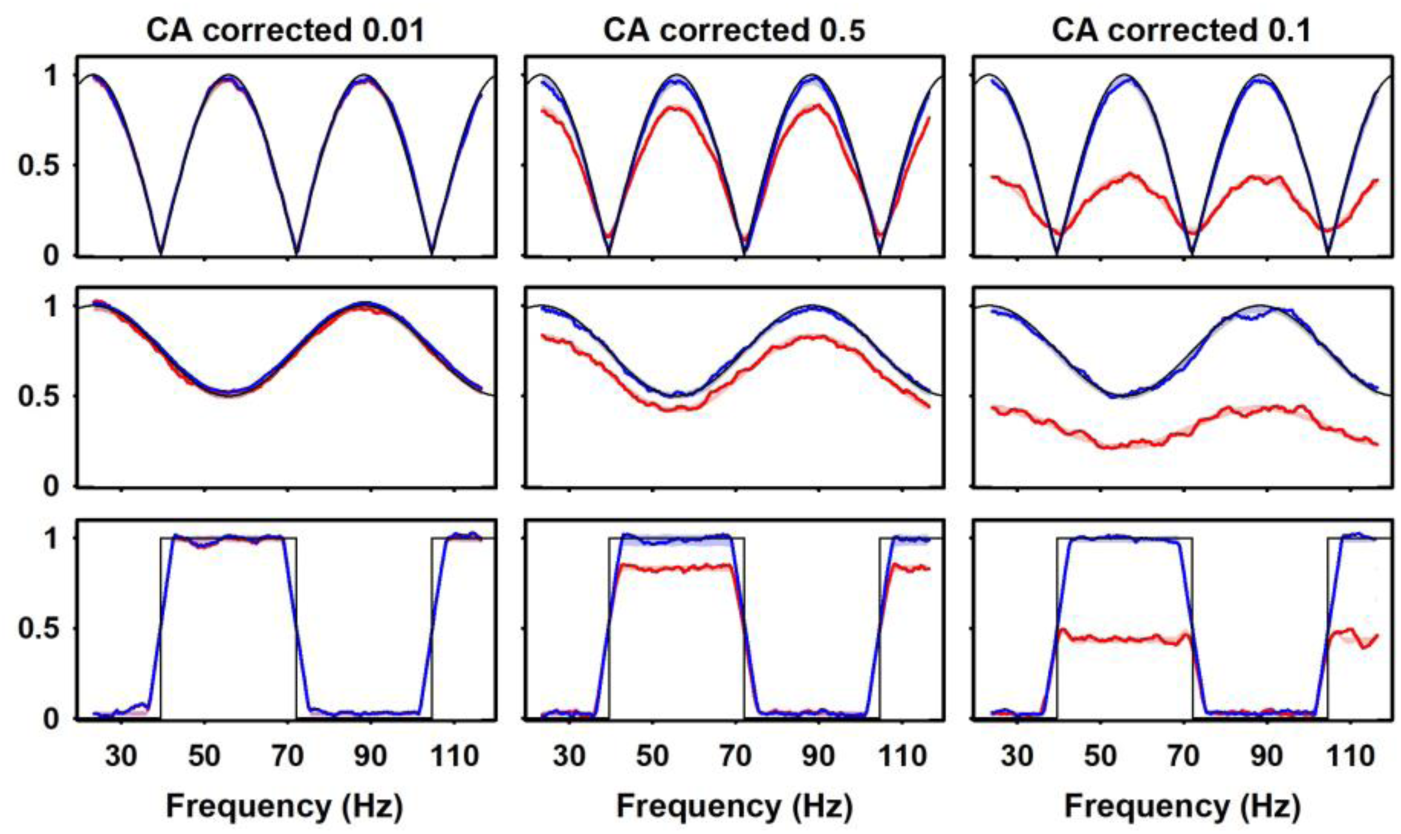
Correcting the EFR obtained by the CA method with the corresponding estimated delay. Simulations were carried out using a pSNR = 2 for the three different EFR shapes (rows) and three true delays (10, 50 and 100 ms, columns). In each panel, the black dots represent the EFR estimated by using the exact true delay (i.e. the best possible performance); the red line represents a single repetition of the EFR estimated without correcting for the delay (i.e. assuming that there is no delay at all, but using this process to estimate the delay) and the blue line represents the same single repetition of the EFR re-estimated using a correction with the delay obtained from the mode of all delays in different IMF. Red and blue shadows represent the area covered by one standard deviation around the mean EFR across all 50 repetitions for the uncorrected and corrected estimation, respectively.

### B. Analysis of real data

In the analysis of electrophysiological recordings in adult rats and human babies, the EFR computed with the three methods, were compared with those obtained with those described in (Prado-Gutiérrez et al. 2012) and (Mijares-Nodarse et al. 2012), respectively, which were obtained using the FA method. Since these authors acquired and processed the EFR using a commercial evoked potential acquisition system MASTER (John et al. 2000) the exact implementation of the FA was not available to us. This impeded us from directly comparing the amplitudes of both estimated EFRs, as they could be differently standardized or given in different units. For instance, the maximum amplitude obtained with the FA in rats was around 0.12 uV (for 70 dB-SPL) and with the STFT was around 0.18 uV, while the CWT and CA showed much higher maximum values over 0.30 uV. In the data of the human baby, these differences were even larger: the maximum value of the FA was almost 0.10 uV, while that for the STFT was of 16 uV and for the CWT and CA this value was over 20 uV

To allow the comparison of the shape of the EFR, we normalized all responses to have the same maximum values as the corresponding EFR obtained with the MASTER system. In general, the shape of the EFR estimated in an adult rat with the three methodologies did not remarkably vary from one to another (Fig. 9). Nevertheless, when the responses were elicited by 70-dB-SPL stimuli, the EFR amplitudes obtained with the STFT and CA in the 140–160 Hz range, seemed to be slightly smaller than those computed with the FA. Moreover, the EFR obtained with the CWT presented higher variability and more local extremes than the responses estimated with the other methods. According to the results obtained with simulated data, this finding can be explained by the higher sensitivity to noise of the CWT. In practice, this effect could be ameliorated by smoothing the response of CWT with a larger time window in a moving average or with by low-pass filtering.

**Fig. 9.**
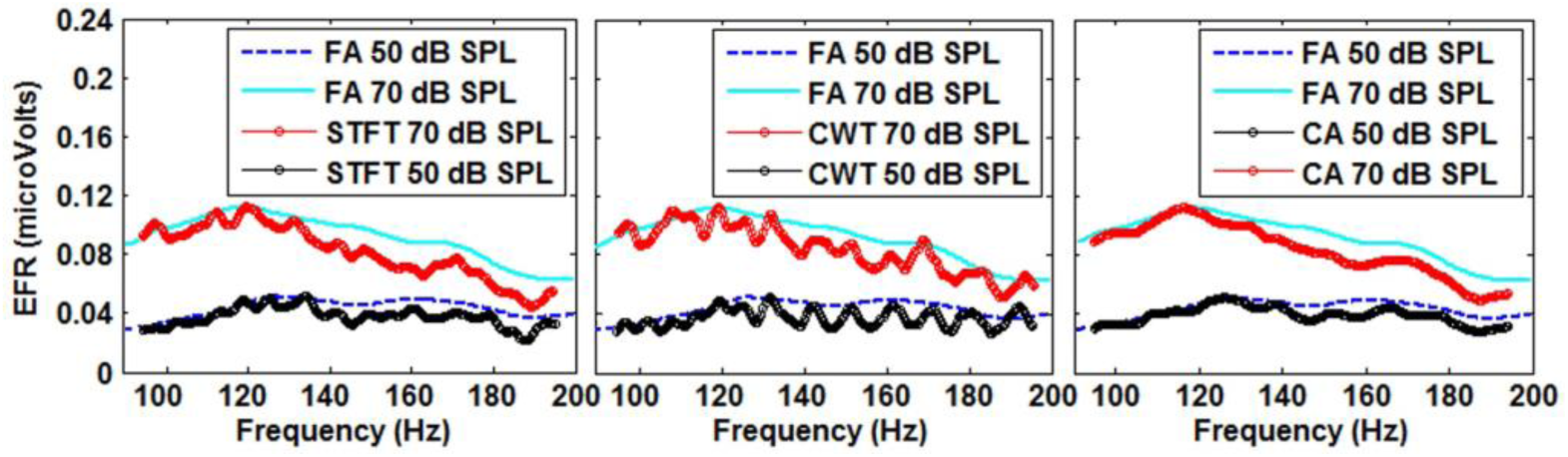
EFRs estimated using the three methods STFT (left), CWT (center) and CA (right), from real electrophysiological recordings of an adult rat stimulated with broadband noise at 50 and 70 db-SPL. The responses were smoothed using a 7-point moving average, and normalized such that all maxima values coincide with those obtained with the FA, for an easier comparison of their shapes. We kept the original amplitudes of the FA responses in microvolts.

Fig. 10, left panel, shows the non-normalized EFR amplitudes of an adult rat, estimated by the three methods. The three methods showed the same optimal modulation frequency (OMF) around 120 Hz for stimulation at 70 dB-SPL and a little higher for stimulation with 50 dB-SPL. The main differences observed are consistent with the results in simulations, namely: the smallest amplitudes corresponded to the evoked potentials obtained with the STFT and the noisiest response was obtained with the CWT. Moreover, the amplitudes of the EFR estimated with the CWT were higher than those obtained with the CA in the whole frequency range. This difference might suggest the presence of a delay between the response and the stimulus, which leads to a decrease in the amplitude of the EFR estimated with CA. Therefore, we estimated the delay as the mode of instantaneous delays obtained with the CA, and then, corrected all EFR estimations using this delay (Fig. 10, right panel). As a result, the only EFR that showed a clear increase in amplitude was the CA, but the OMF and the general shape of the response was kept. In this case the estimated delay was about 24 ms, using both the mode and median.

**Fig. 10.**
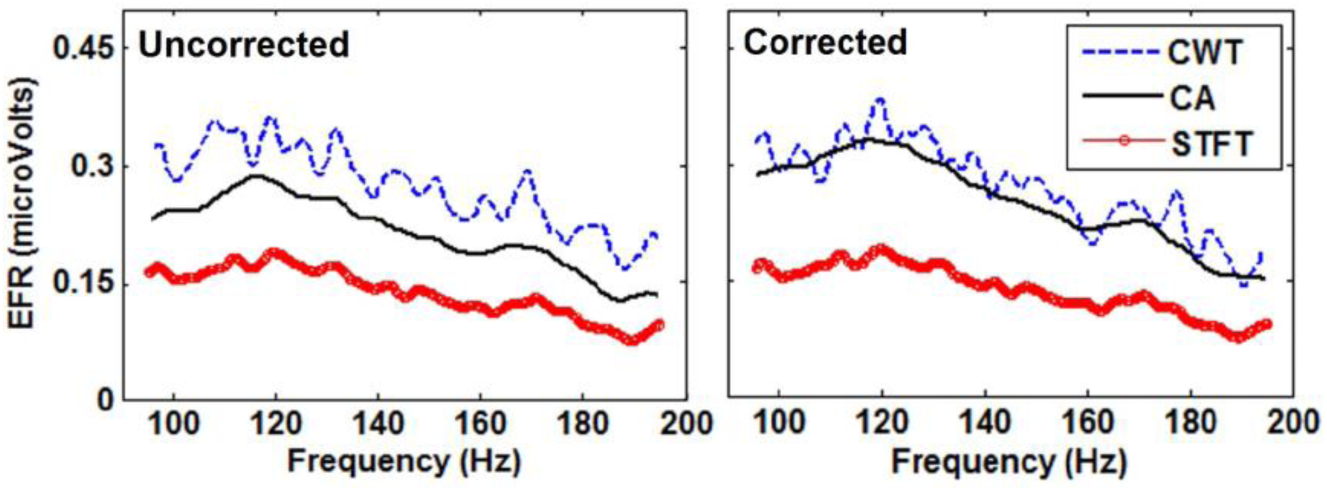
Left: Non-normalized EFRs estimated with the three methods studied, from a real electrophysiological recording in an adult rat stimulated with broadband noise at 70 db-SPL. Right: the EFRs obtained for the same data as in the left panel but correcting the estimation with the delay estimated with the mode of instantaneous delays measured with the CA method.

Fig. 11 shows the same comparison as in Fig. 9 but for a real data recorded from a human baby. Like the results found in rats, the shape of estimated EFRs with the three methods mimic that of the FA, with the OMF being very similar in all cases. However, the difference in EFR amplitudes obtained for frequencies higher than 110 Hz are more consistent and slightly larger than in rats, especially for the STFT and the CA methods. The estimated EFRs without normalization, presented also the same general behavior as for the data from rats. Fig. 12, left panel, shows that STFT offered the smallest amplitudes and the CWT the highest and noisiest. Fig. 12, right panel, shows the EFRs re-calculated using the delay estimated as the mode of instantaneous delays given by the CA method. Again, only the EFR estimated with the CA showed a noticeable increase in the amplitude. Indeed, the values of the delay of the response were around 34 ms, which is large enough to make the CA underestimate the actual EFR amplitudes.

**Fig. 11.**
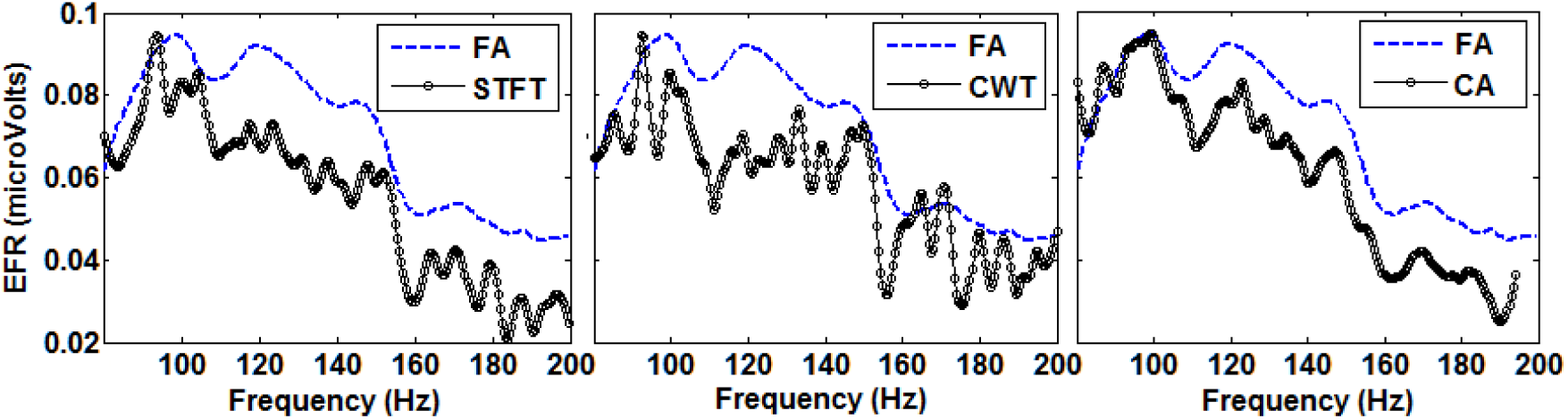
EFRs estimated using the three methods STFT (left), CWT (center) and CA (right), from real electrophysiological recordings of a human baby stimulated with broadband noise at 50 db-HL. The responses were smoothed using a 7-point moving average, and normalized such that all maxima values coincide with those obtained with the FA, for an easier comparison of their shapes. We kept the original amplitudes of the FA responses in microvolts.

**Fig. 12.**
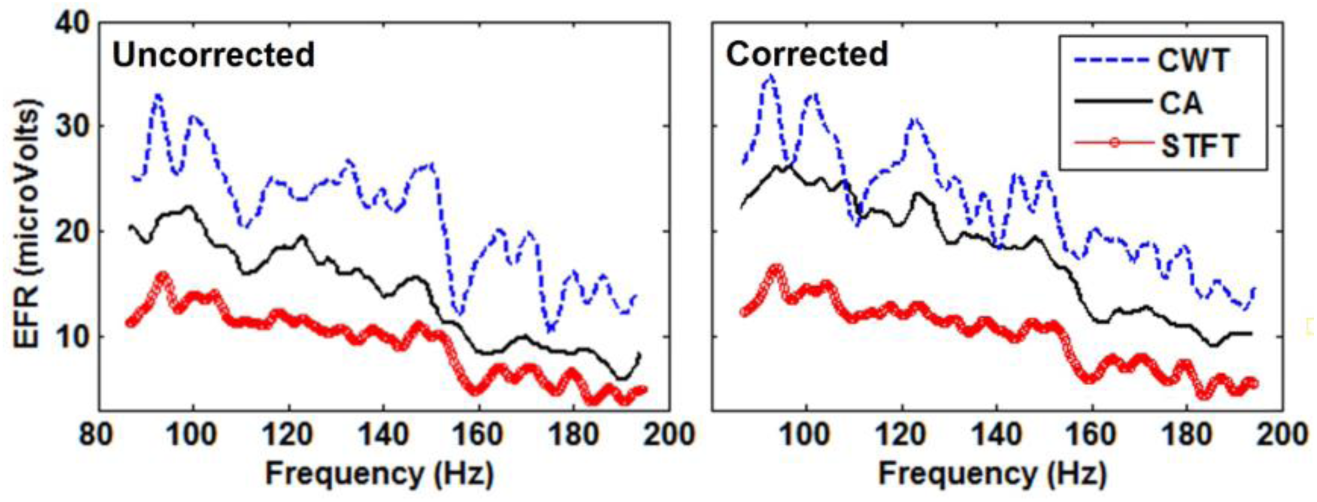
Left: non-normalized EFRs estimated with the three methods studied, from a real electrophysiological recording in a human newborn stimulated with broadband noise at 50 db-HL. Right: non-normalized EFRs after correcting the estimation by using a delay of the response with respect to the stimulus. This delay was estimated as the mode of all instantaneous delays computed with the CA.

## IV. Discussion

In this work, we have introduced a new method for estimating auditory envelope following responses (EFR). This method is based on using the amplitude envelope of the AM-stimulus as reference function for a sliding-window correlation with the recorded EEG. As the EFR is commonly elicited by synthetic acoustic stimuli in which the amplitude is modulated with a linear chirp, we have called the method Chirp Analyzer (CA). The CA is very similar to the Fourier Analyzer, but better copes with the non-stationarity of the electrophysiological responses to those stimuli with ever-changing frequency. This type of response has been studied in different experimental conditions and different methodologies and all studies support our main assumption that the response will instantaneously follow the modulation signal of the stimulus (Artieda et al. 2004; Aiken & Picton 2006). In addition, the method proposed here can theoretically be used in the analysis of any other brain response (visual, or somatosensory) that follows a continuous stimulation with non-stationary characteristics, or more generally, where the time-frequency properties of the response are known.

### A. Evaluating the performance of time-frequency methods with simulations

Simulations showed that the CA is more robust and reliable than the STFT and the CWT to estimate EFR with different shapes, even in presence of noise (Fig. 4 and 5). Importantly, the EFR estimated with the STFT method, which is equivalent to the FA, is much more biased towards lower amplitudes those of the other methods. This is expected given that the STFT uses stationary reference signals that can never match the response in the entire windows of analysis, thus intrinsically estimating lower amplitudes due to its spectral underrepresentation. This bias might not be so important when the interest is in characterizing the relative amplitudes of the EFR, i.e., determining the OMF of the response or those frequencies with particularly low responses, but it makes the STFT and FA methods not reliable to estimate actual absolute amplitudes of the response, which might be relevant for its proper statistical detection.

The CA provided good estimates for very low response amplitudes with higher robustness than the CWT method. The latter showed to be the most sensitive to noise, giving a nonzero estimation of the EFR amplitude even for those frequencies where there was no response at all. This is a consequence of the fixed proportionality between frequency and temporal resolution implemented in the CWT to obtain an adequate tradeoff between time and frequency resolution. In the case of non-stationary signals like EFR, the optimal proportionality constant is difficult to assess and strongly depends on the range of frequencies covered by the stimuli. In our study, the wide frequency range (20 to 120 Hz) lead to a very high temporal resolution for higher frequencies, making the EFR estimation to be much noisier than those offered by the STFT and the CA. As a positive point, the high temporal resolution allows to estimate abrupt changes in amplitude better than the other two methods and to offer unbiased amplitudes in averaged signals, although with higher variance.

The proposed CA method showed the worst performance in terms of robustness to the presence of a temporal delay in the EFR, i.e., in cases where the responses followed the stimulus with a delay that may have physiological (latency of the response) and/or instrumental origin (Fig. 6). This was not surprising as the CA is based on correlating the recorded signals with the reference chirp assuming a zero delay, while STFT and CWT estimate a whole time-frequency energy that may expand to neighbor frequencies depending on the spectral resolution. As the modulation is non-stationary, any displacement between the signal and the reference chirp will make them to have several cycles in counter-phase, leading to a quick drop in the correlation. However, we found that for delays lower than 50 ms the underestimation of the EFR amplitude given by the CA is smaller than the typical bias showed by the STFT estimation (even in the absence of a delay), and the CA is still able to recover the correct shape of the response in the whole range of modulation frequencies (Fig. 6).

Several strategies can be followed in order to minimize the influence of the unknown latency of the response. On one hand, the experimental stimuli should use modulating linear chirps with small slope for the linear change of frequency with time, and the analysis window should also be large enough to minimize the influence of transient activities. In this sense, Aiken & Picton (2006) showed that using slopes smaller than 10 Hz/s the evokes responses can follow the stimulation and can be considered locally stationary for the analysis with the classical Fourier Analyzer. On the other hand, a more direct approach would be to introduce the knowledge about the latency of the response in the procedure for estimating the EFR. However, correct values of delays can vary in every experiment and in every individual. The alternative is to find a reliable way to estimate the delay between stimulation and response that can adapt to specific experimental conditions and brain responses.

### B. Evaluating the estimation of response delays

The estimation of the actual delay between stimulus and measured responses in this kind of continuous repetitive stimulation experiments (usually known as steady-state responses), has been attempted in previous studies using a linear regression between the phase difference (between stimulus and recorded signal) and the modulation frequency (Kuwada et al. 2002; Pauli-Magnus et al. 2007; Prado-Gutierrez et al. 2012). The delay can be computed from its proportional relation with the regression slope. However, this method is very sensitive to the correct estimation of phases in real signals. The circular nature of phases makes this estimation problematic as several manipulations are needed to convert them in a linear magnitude that can be subject to regression, and these manipulations do not always ensure the correct unwrapping of the phases. Moreover, recent studies have proven that when the amplitude of oscillatory responses is low, the estimation of phases using classical time-frequency methods based on Fourier basis, are not reliable and thus any computation using them would not be trustable (Sameni & Seraj 2017).

In our simulations, we found that applying this methodology with the STFT and CWT did not provide correct estimates of the delay, since the phases were consistently miss-estimated in most of the simulated scenarios (see Tables I and II). The CA showed better estimates of the delay when using the regression but always with errors over 10 ms. We also confirmed that these problems for estimating the phases worsen when the simulated EFRs had very low amplitudes in specific modulation frequencies (Fig. 7).

The new method for estimating delays proposed here are based on finding the time between the stimulation and the maximum amplitude of the response, for each IMF of the stimulus. Although this approach allows the estimation of one delay for each IMF, we assume that the physiological delay is the same for all of them except for statistically random variations, given that the stimulus properties change slowly with time. Therefore, the response delay can be simply computed by using population measures over the whole set of instantaneous delays, namely the mean, the weighted mean, the median and the mode.

Results presented in Tables I and II showed that the STFT and the CA can be used for reliably estimating delays, while the CWT is not recommended for estimating the response delay due to its higher sensitivity to noise. Since the instantaneous delays computed with the STFT showed a Gaussian distribution around the true delay (except in those IMFs where the EFR amplitude was too small) the use of population measures was appropriate in this case. However, in the case of the CA, the distribution of the instantaneous delays was not Gaussian, as these values were mostly underestimating the true delay. For this method, it is more convenient to use the mode across all instantaneous estimated delays (see Tables I and II and Fig. 7). The mode also has the advantage of being a value that belongs to the population, which in this case is an integer number of time steps. Therefore, using the mode as the descriptor of the population makes the estimation of the delay more robust to outliers and other wrong estimations. For the CA, even in the worst signal-to-noise ratio scenario, only 8 out of the 50 repetitions showed the mode of instantaneous delays to be different from the true delay.

However, an important practical limitation here is that these results depended on the overlapping of the analysis windows for the estimation of instantaneous delays, which at the same time also determines the computational time of the method. A high overlapping will be more likely to lead to the true delay but computation could take up to 1 minute (for a single repetition). In our data, we found that for a time step between consecutive windows of up to 100 ms (overlapping of 90% for one-second-long windows), the mode can still give the true delay. Nevertheless, for higher values (up to 200 ms), the absolute errors can be of about 8 to 15 ms, although the algorithm will run in only 5 seconds. This means that the CA could be used to estimate the delay only when small delays are expected, following the strategy of using a high temporal overlapping only for estimating the delay once. The estimated delay can then be used for correcting the estimation of the EFRs with a lower temporal overlapping. Alternatively, when delays are expected to be higher than 50 ms, we recommend the use of the STFT for estimating a reliable value of the delay with the weighted mean or the median of the instantaneous delays.

### C. Correcting the EFR by the estimated delay

When the EFR was estimated without knowledge of a nonzero response delay, only the CA was sensitive enough to show a decrease of about 20% in the amplitudes, for a delay of 50 ms and of more than 50% when the delay was 100 ms. However, when delays were around 10 to 20 ms, the difference between the estimated EFR without considering the delay and the true simulated EFR was less than 2% (relative error). This relative error of around 1 to 2% was also found when comparing the corrected EFR using the true delay and the uncorrected EFRs when the delay was 10 ms (Fig. 8, left column). Therefore, these results suggest that if we are mainly interested in estimating the EFR and delays are expected to be around 10 ms or less, there is no real need to compute a new EFR corrected by the estimated delay.

On the other hand, if delays are expected to be higher than 40 or 50 ms, then the correction ensures a much better reconstruction of the EFR. In those cases, using small time steps would be necessary for a reliable estimation of the delay with the CA. In our simulations, we used full overlapping windows for estimating the delay with the CA (i.e. subsequent windows overlap in all but one time point, which corresponds to a time step of 0.5 ms), with the estimation taking about 50 seconds for the analysis of one repetition. However, as mentioned before, we found that there were no large errors in the estimation of the delay if the time step is increased up to 50 ms, with the estimation taking only around 5 seconds for one repetition. When comparing the corrected EFR using the estimated delays with another corrected EFR using the true simulated delay, we found that in all cases, the relative error was lower than 2%, even when the estimated delays were different from the true delay in up to 10 ms. Specifically, if the delay is estimated using the mode, weighted mean or median (the last two cases always led to underestimating the true delay in a few milliseconds) the relative error was below 1%. Only when using the mean as the population measure, the relative error reached 2% in a few repetitions, mostly in the case of the Rect-deep scenario, where there are many IMFs without response (Fig. 8).

As discussed above, a higher precision in the estimation of the delay could be obtained by using the STFT. Nevertheless, given that this method is around 20 times slower than the CA, we consider unpractical using it for estimating latencies in clinical settings. If the objective is to measure physiological delays in well-controlled experiments, then the STFT showed to be the method of choice for large delays.

### D. Physiological limitations for a proper estimation of the response delay

Despite the methodological limitations discussed in previous subsection for estimating a reliable response delay in practical experiments, there are factors that depend on the specific experiment, and even on the individuality of the experimental subject. The most relevant is the possibility of finding individuals with pathological responses or using specific stimuli that do not evoke response in some of the instantaneous modulation frequencies (IMFs) used. If the responses are very small in a particular range of IMFs, this will lead to less reliable estimations of both the EFR (usually overestimated amplitudes) and the instantaneous delays, which will influence the final estimation of the response delay.

Physiologically, the lack of response to a specific range of IMFs might reflect different sensitivities to the intensity of the stimuli or changes with age in the behavior of the cochlea or the external auditory system, which usually is of clinical relevance. Therefore, in future studies, it would be important to establish the conditions under which the methods studied here would be useful for reliably characterizing responses from these individuals. As a very preliminary exploration, we prepared a set of simulations using an EFR with the Rect-deep shape, but only with one rectangular range of frequencies with unit amplitude and zero elsewhere. The width of the range of IMFs (hereinafter the “bandwidth” of the response) was varied, as a direct way of simulating the range of frequencies where envelope-following responses are high (i.e. varying the amount of “good” data to estimate both EFR amplitude and delays). This allowed us to study what is the minimum number of IMFs (that evoke high-amplitude responses) needed to trust the estimated delay with the proposed methods.

Fig. 13 shows the estimated delays using the STFT and CA with different measures, with respect to the bandwidth of the simulated response. In accordance to results presented above (Fig. 7), using the STFT method, both the weighted mean and the median provided robust estimations of the delay, which was similar to the true delay when the bandwidth was around 15 Hz, although stabilize on the true delay when the bandwidth reaches 60 Hz (Fig. 13, left). The good performance for relatively narrow bandwidth (15 Hz) is explained by the gaussian nature of instantaneous delays estimated with the STFT around the true delay, such that a few data points are enough for a rough estimation. However, it needs more data points for a perfect estimation than the CA method, which showed higher variability for small bandwidths, but the mode was able to estimate the exact true delay from bandwidths of 17 Hz or more, and the median from around 23 Hz (Fig. 13, right). The mode in the case of the STFT was close but never perfect and the method based on a linear regression between phase difference and IMFs, needed too many data points (almost the full 100Hz bandwidth) for a good estimation in this specific simulation. In practice, this exploration suggest that the methods are also robust for the lack of data and can be used in a wide range of normal and pathological cases. In those cases, a recommendation could be to firstly estimate an EFR to evaluate if there are enough IMFs with high amplitudes that can support a good estimation of the response delay for a subsequent correction of the EFR.

**Fig. 13.**
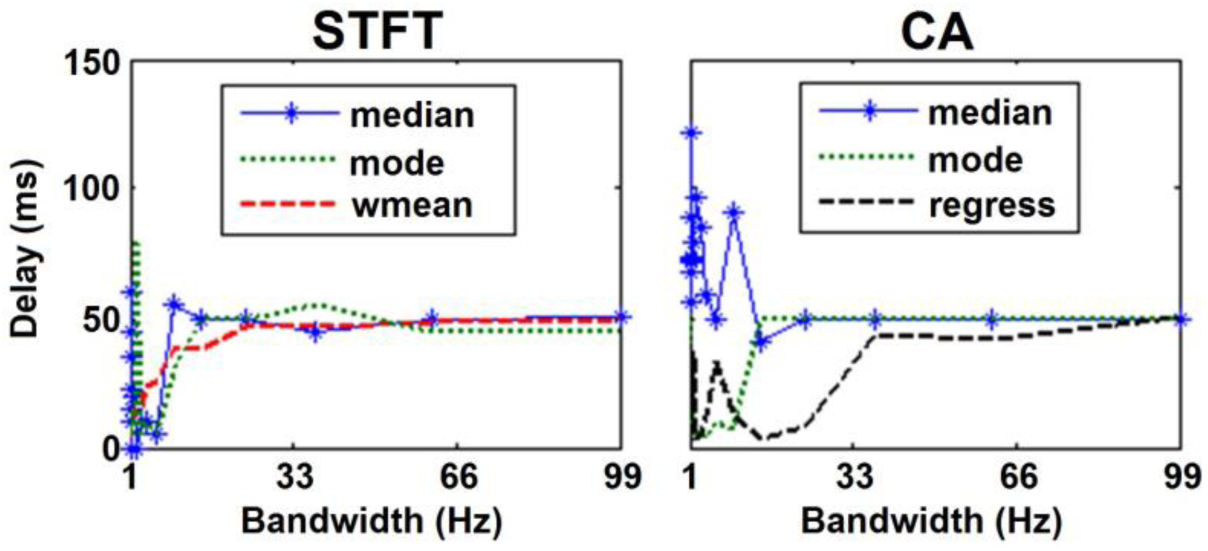
Delay estimated by the STFT (left panel) and CA (right panel) with different measures, from simulated signals using a rectangular pulse-like EFR with varying width of the range of frequencies showing nonzero (unit) amplitudes (Bandwidth). The signals were simulated with a true delay of 50 ms and pSNR=2.

### E. Analysis of EEG recordings in rats and human newborns

#### 1) Estimation of the amplitude of the EFR

In this work, the analyses of real data allowed us to make a preliminary evaluation of the proposed methods, in comparison with the FA method available in the commercial system MASTER. In general, the EFR estimated with the STFT, CWT and CA showed the same general shape with respect to modulation frequencies and similar OMF, as compared to the results found with the FA method, which has been the most popular method for estimating these type of auditory responses (Purcell et al. 2004; Aiken & Picton 2006; Prado-Gutiérrez et al. 2012; Mijares-Nodarse et al. 2012). In both datasets, the behavior of estimated EFR for the different methods was also consistent with our findings in the analysis of simulated data. Although we believe that a thorough validation should be carried out with larger datasets, our results suggest that the methods explored here are promising tools for a proper characterization of the experimental EFR.

However, two main differences appeared when comparing the EFR estimated with the FA and those estimated with the time-frequency methods and the CA: the absolute amplitude of the EFR in every IMF and the level of smoothness for the whole curve. The differences in amplitudes might be explained by using different normalization in the signal processing procedures, which is a common issue in this type of real data analysis. In practice, this issue is ignored as the absolute values are not the main interest but just the relative amplitudes for different modulation frequencies. However, according to our simulations, we were indeed able to recover the right amplitude with the CWT and CA methods, while the STFT (equivalent to FA) was consistently biased toward lower amplitudes. This might also explain why the FA method offered lower absolute amplitudes of the EFR. The differences in the smoothness of the EFR are more difficult to explain given that the original estimation of the EFR with the FA was not available. It might be possible that the FA have been smoothed in a stronger way than the simple moving-average 7-point smoother applied to the responses estimated with the other methods. Of course, it is hard to know what is the suitable level of smoothing in the case of real data analysis, thus we decided not to use a strong smoothing in order to evaluate variability and local maxima. It is possible that increasing the level of smoothing might make all curves to look much more similar.

Other explanations for the differences found may lie in the limitations of the proposed methods, according to the assumptions made on the real unknown nature of the physiological response. From a physiological point of view, it could be possible that the activity of neural generators of the EFR does not follow exactly the model assumed. For instance, the biological system might not respond continuously to every tiny change in the modulation frequency, but instead it responds preferentially to specific frequencies by keeping the frequency of the response fixed for a range of different IMFs in the stimulus. Although this behavior corresponds to a piece-wise stationary response, the exact division of stationary segments would not be known a priori. In that scenario, time-frequency methods still seem a better option than FA, since they allow the exploration of the changes in the signal before selecting ad hoc the relevant time-frequency pairs where to extract the estimated EFR. Finally, it should be recognized that the electrical activity recorded with the EEG reflects the sum of the activity in many different brain sources that may show non-linear behavior at the level of neuronal masses, which are difficult to disambiguate from the macroscopic signal. In this sense, future research should include the design of specific experiments to study this type of physiological response at the neuronal level.

#### 2) Estimation of the latency of the EFR

The estimation of the delay of the response in real data is also a promising contribution of this work. In the analysis of the real data from both rats and human babies, we found that only the EFR estimated by the CA showed an increase in amplitude when it was corrected by the delay, which suggest that there was indeed an actual nonzero delay between the stimulus and the recorded signal. Interestingly, the delays estimated in the real datasets analyzed here did not coincide with values usually handled in the literature. In rats, we obtained delays in the order of 24 ms, while the delays reported in the literature for EFR generated in subcortical structures are below 5 ms (Prado-Gutierrez et al. 2012). For the human newborn, the estimated delay around 34 ms is also higher than the other estimations of the apparent delay that has been reported to be between 9 to 21 ms according to different frequency ranges (Purcell et al. 2004). Despite the possibility of having contributions to the delay of the measured activity given by other non-physiological sources, we might argue on a few other factors that may explain these differences. Firstly, most of the reported delays in the literature have been found using the regression method, which heavily relies on the correct estimate of phases of the oscillatory responses. The correct estimation of phases is not straightforward, and several studies have shown the potential misleading results and misinterpretation when using methods based on mathematical measures of estimated phases in real signals (Sameni & Seraj 2017, Martinez-Montes et al. 2008). Moreover, phase differences are undetermined by an integer number of full 2π-cycles. Secondly, the noisy nature of the signals might also influence this estimation with any method. The level of noise in the real data (especially from human newborns) is probably higher than the one used in our simulations, and it is possible that the population measures we used here (mean, weighted mean, median, mode) are not appropriate for a reliable estimation of these delays in noisier data, especially if they are smaller than 10 ms. In any case, these discrepancies need to be confirmed in future studies with larger dataset, and preferably with novel experiments especially designed for this purpose.

## V. Conclusions

In this work, we have formulated the estimation of the Envelope Following Response (EFR) with the use of explicit time-frequency methods, which offer more information about the energy distribution of the recorded signal than the traditionally used Fourier Analyzer (FA) method. Importantly, we further introduced the Chirp Analyzer (CA) method as a new tool for the estimation of the EFR. Instead of using a Fourier basis as reference signals for the estimation of the EFR, the CA uses the same linear chirp that modulates a carrier tone in the stimulus as the reference function. If the linear change in instantaneous modulation frequency is small enough, the response can be considered a steady-state response and it will closely follow the chirp modulation function. Therefore, the CA allowed for a better match between the recorded signal and the reference function than the FA, which directly impact the estimation of the response parameters.

In a direct comparison using controlled simulated responses, the CA showed to be able to estimate the correct EFR amplitude, without the typical bias obtained when using the Short-Term Fourier Transform (equivalent to FA but for the whole time-frequency amplitude). It was also more robust to noise than the Morlet Continuous Wavelet Transform. However, CA is more sensitive to the presence of a delay in the response with respect to the stimulus; therefore, it should be used cautiously when this delay is expected to be higher than 50 ms. For coping with this last issue, we proposed here a novel methodology for estimating the apparent latency of the response. This method proved to be reliable when using the STFT and the CA methods, as assessed using simulated responses. The estimation of the EFR amplitude with any of the methods, but especially with CA, should be corrected by using the estimated delay when possible.

Analysis of real data suggested that all time-frequency methods were able to estimate EFR amplitudes, but they should be interpreted in the light of the limitations shown in the simulation studies. Finally, we hope that the results obtained in this study contribute to improve and standardize the tools for the analysis of the EFR that are currently available. This is important for the future extended use of this kind of response in audiology. The strategy followed here for introducing the CA is very general and can be applied to almost all types of oscillatory brain responses, therefore, it opens the possibility of more complex experiments that evaluates brain responses to stimulus with properties varying in a wide range in a single run.

## References

1. Aiken SJ, Picton TW (2006) Envelope following responses to natural vowels. Audiol Neurotol 11: 213–232.

2. Artieda J, Valencia M, Alegre M, Olaziregi O, Urrestarazu E, Iriarte J (2004) Potentials evoked by chirp-modulated tones: a new technique to evaluate oscillatory activity in the auditory pathway. Clin Neurophysiol 115: 699–709.

3. Boashash, B (2003) Time frequency signal analysis and processing. Elsevier, London.

4. Choi, JM, Purcell DW, Coyne JAM, Aiken SJ (2013) Envelope following responses elicited by English sentences. Ear and hearing 34, 5: 637–650.

5. Dajani H, Purcell D, Wong W, Kunov H, Picton TW (2005) Recording human evoked potentials that follow the pitch contour of a natural vowel. IEEE Trans Biomed Eng 52: 1614–1618.

6. Durka, P. (2007) Matching pursuit and unification in EEG analysis. ArtechHouse, London.

7. Gurtubay IG, Alegre M, Labarga A, Malanda A, Iriarte J, Artieda J (2001) Gamma band activity in an auditory oddball paradigm studied with the wavelet transform. Clin Neurophysiol 112: 1219–1228

8. Kiebel, S. J., Tallon-Baudry, C., & Friston, K. J. (2005). Parametric analysis of oscillatory activity as measured with EEG/MEG. Human brain mapping, 26(3), 170–177.

9. Kuwada S, Anderson JS, Batra R, Fitzpatrick DC, Teissier N (2002) Sources of the scalp-recorded amplitude-modulation following response. J Am AcadAudiol 13: 188–204.

10. Laroche, M., Dajani, H. R., Prévost, F., & Marcoux, A. M. (2013). Brainstem auditory responses to resolved and unresolved harmonics of a synthetic vowel in quiet and noise. Ear and hearing, 34(1), 63–74.

11. Martínez-Montes, E., Cuspineda-Bravo, E. R., El-Deredy, W., Sánchez-Bornot, J. M., Lage-Castellanos, A., Valdés-Sosa, P. A. (2008). Exploring event-related brain dynamics with tests on complex valued time–frequency representations. Statistics in medicine, 27(15): 2922–2947.

12. Mijares-Nodarse, E., Pérez Abalo, M. C., Torres Fortuny, A., Vega Hernández, M., & Lage Castellanos, A. (2012). Maturational changes in the human envelope-following responses. Acta Otorrinolaringologica (English Edition), 63(4), 258–264.

13. Pauli-Magnus D, Hoch G, Strenzke N, Anderson S, Jentsch TJ (2007) Detection and differentiation of sensorineural hearing loss in mice using auditory steady-state responses and transient auditory brainstem responses. Neuroscience 149: 673–684.

14. Perez-Alcazar M, Nicolas MJ, Valencia M, Alegre M, Iriarte J, Artieda J (2008) Chirp-evoked potentials in the awake and anesthetized rat. A procedure to assess changes in cortical oscillatory activity. Exp Neurol 210: 144–153.

15. Picton TW, Dimitrijevic A, John MS (2002) Multiple auditory steady-state responses. Ann Otol Rhinol Laryngol Suppl 189: 16–21.

16. Prado-Gutierrez P, Mijares-Nodarse E, Savio G, Borrego M, Martínez-Montes E, Torres A (2012) Maturational time course of the Envelope Following Response to amplitude-modulated acoustic signals in rats. International Journal of Audiology 51(4): 309–316.

17. Purcell DW, John MS, Schneider BA, Picton TW (2004) Human temporal auditory acuity as assessed by envelope following responses. J Acoust Soc Am 116(6): 3581–3593.

18. Purcell DW, John MS (2010) Evaluating the modulation transfer function of auditory steady state responses in the 65 Hz to 120 Hz range. Ear Hear 31: 667–678.

19. Rickards FW, Tan LE, Cohen LT, Wilson OJ, Drew JH, Clark GM (1994). Auditory steady-state evoked potentials in newborns. British Journal of Audiology, 28: 327–337.

20. Regan, D. (1989). Human Brain Electrophysiology: Evoked Potentials and Evoked Magnetic Fields in Science and Medicine, New York: Elsevier Science, pp. 70–98, 112 – 123, 273 – 275.

21. Sameni, R., Seraj, E. (2017). A robust statistical framework for instantaneous electroencephalogram phase and frequency estimation and analysis. Physiological measurement, 38(12): 2141.

